# CASowary: CRISPR-Cas13 guide RNA predictor for transcript depletion

**DOI:** 10.1101/2021.07.26.453663

**Authors:** Alexander Krohannon, Mansi Srivastava, Simone Rauch, Rajneesh Srivastava, Bryan C. Dickinson, Sarath Chandra Janga

**Author notes:** Correspondence should be addressed to: Sarath Chandra Janga, 719 Indiana Avenue Ste 319, Walker Plaza Building, Indianapolis, Indiana – 46202.

## Abstract

Recent discovery of the gene editing system -CRISPR (Clustered Regularly Interspersed Short Palindromic Repeats) associated proteins (Cas), has resulted in its widespread use for improved understanding of a variety of biological systems. Cas13, a lesser studied Cas protein, has been repurposed to allow for efficient and precise editing of RNA molecules. The Cas13 system utilizes base complementarity between a crRNA/sgRNA (crispr RNA or single guide RNA) and a target RNA transcript, to preferentially bind to only the target transcript. Unlike targeting the upstream regulatory regions of protein coding genes on the genome, the transcriptome is significantly more redundant, leading to many transcripts having wide stretches of identical nucleotide sequences. Transcripts also exhibit complex three-dimensional structures and interact with an array of RBPs (RNA Binding Proteins), both of which further limit the scope of effective target sequences. As a result, there currently exists no method to predict whether a specific sgRNA will effectively knockdown a transcript. Here we present a novel machine learning and computational tool, CASowary, to predict the efficacy of a sgRNA. We used publicly available RNA knockdown data from Cas13 characterization experiments for 555 sgRNAs targeting the transcriptome in HEK293 cells, in conjunction with transcriptome-wide protein occupancy information on RNA. Our model utilizes a Decision Tree architecture with a set of 112 sequence and target availability features, to classify sgRNA efficacy into one of four classes, based upon expected level of target transcript knockdown. After accounting for noise in the training data set, the noise-normalized accuracy exceeds 70%. Additionally, highly effective sgRNA predictions have been experimentally validated using an independent RNA targeting Cas system -CIRTS, confirming the robustness and reproducibility of our model’s sgRNA predictions. Utilizing transcriptome wide protein occupancy map generated using POP-seq in Hela cells against publicly available protein-RNA interaction map in Hek293 cells, we show that CASowary can predict high quality guides for numerous transcripts in a cell line specific manner. Application of CASowary to whole transcriptomes should enable rapid deployment of CRISPR/Cas13 systems, facilitating the development of therapeutic interventions linked with aberrations in RNA regulatory processes.

## Background

Gene editing technologies have played an increasingly important role in numerous life science domains in the recent years, especially in the fields of biology, biotechnology, and medicine (1). At the center of many of these discoveries is the CRISPR/Cas9 gene editing system (2). This system has allowed an unprecedented level of accurate and precise editing of the genome. Several limitations have been recognized with the use of CRISPR/Cas9 system: the requirement of a PAM (protospacer adjacent motif) sequence adjacent to the target gene sequence, reliance on dynamic DNA repair procedures (3), and its inability to facilitate tissue specific alterations (4). However, other Cas proteins are being identified and repurposed as systems for genome and transcriptome editing (5).

One such class of protein, Cas13, has been modified to directly edit RNA transcripts (5). Much like Cas9, the Cas13 system is a two-component system: the Cas13 enzyme and sgRNA. After binding to the sgRNA, the Cas13 complex probes the cellular RNA molecules for a sequence complementary to the 28-nucleotide spacer sequence of the bound sgRNA. Once identified, the enzyme binds to the RNA molecule for its catalytic cleavage, rendering it ineffective and facilitating RNA degradation. Some of the most promising aspects of this system are the independence from the PAM motif restriction and the potential for designing guide sequences for enabling tissue specific transcript knockdowns.

While the Cas13 system does offer some distinct advantages over the Cas9 system, it also poses some unique challenges. First and foremost, most of the transcriptome remains unknown, owing to poor understanding of various post transcriptional processes. RNA molecules can also adopt a variety of complex three-dimensional structures through networks of intramolecular interactions. Additionally, there are a variety of proteins, RNA binding proteins (RBPs), which bind to various regions of RNA molecules. Together the irregular complex structure of RNAs and RBP binding act to limit the number of stretches available for complementary base pairing. Therefore, a tool to predict the efficacy of a given sgRNA is desirable.

To that end CASowary was developed as a first of its kind sgRNA efficacy predictor. Although several previous studies have focused on creating software to predict sgRNA for CRISPR Cas9, to our knowledge, there have been no previous attempts for doing such for CRISPR Cas13 (6–8). CASowary was written in python3 and uses a variety of functions from various libraries: vector operations form numpy, statistical analysis from scipy, machine learning utilities from sklearn, and data visualization from seaborn and matplotlib (9–13). The development and validation of CASowary took place over three distinct phases: Data Collection and Integration, Feature Selection, and Model Generation and Benchmarking (Fig 1). Three different types of data were utilized by the model for predictions -targeted RNA knockdown experiments (14), transcriptome-wide protein occupancy information (15), and sgRNA spacer sequence alignment data. Feature selection took place through a variety of steps including composition analysis, k-mer capture, and evaluating feature significance and contribution. The model was validated using both 3-fold and 5-fold cross-validation. Additionally, the model’s predictions were verified through an experimental protocol with an orthogonal CRISPR based system (16). The model was then applied to all transcripts from among 5900 random genes, to determine any biological relevance of the model’s predictions.

**Figure 1.**
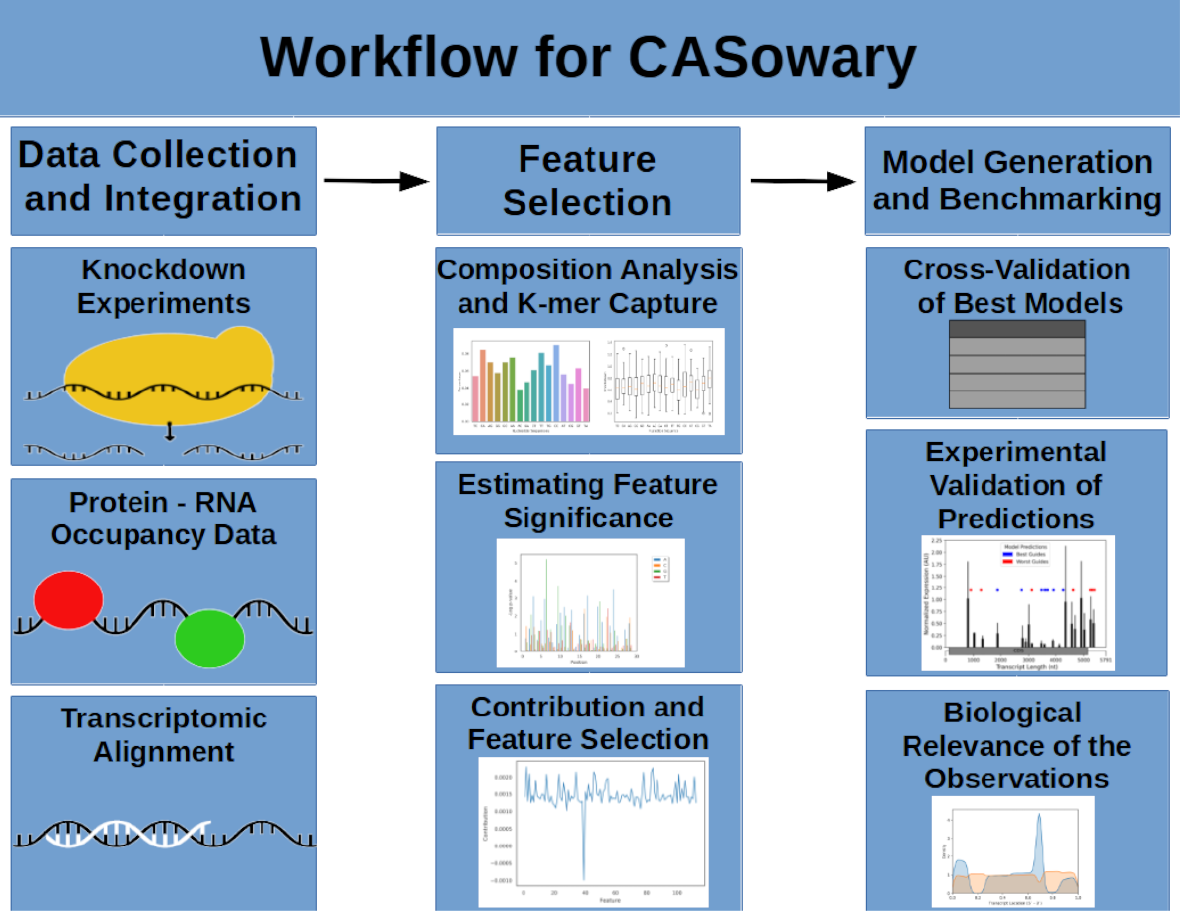
Algorithmic Framework for CRISPR Cas13 Guide RNA Prediction. CRISPR Cas13 knockdown experiments, protein occupancy, and transcriptomic alignment data was gathered for consideration and analysis by the model. Feature lists were created through composition analysis and k-mer capture. The significance and contribution of each feature was estimated to create finalized possible lists of features. The final feature list for the model was generated through comparison of 3-fold and 5-fold cross-validation experiments. Model predictions were validated through direct comparison with performed experiments. The model used to predict sgRNA’s spanning all transcripts associated with 5900 genes. The results were collected and analyzed for any potential biological relevance for predictions.

## Implementation

### Genome-scale sequence data for CRISPR-Cas13

Utilizing the Cas13 human transcript knockdown experiments from Abudayyeh et al (14), we sought to develop a machine learning model that predicts the effectiveness of a given sgRNA at knocking down a target transcript. Firstly, we investigated the sequence composition i.e. mono-, di-, and tri-nucleotide compositions for all 555 guide-RNAs (or sgRNA) at each position along the 28-nt length. We obtained a list of over and under-represented k-mers (chi-squared test) at each location across the spacer sequence of the sgRNA (Fig 2 A-B). Afterwards, sgRNAs were then partitioned into distinct groups based upon their nucleotide composition at a specific location; in order to perform a Kruskal Wallis (17) test. Sets of positions with Kruskal values less than 0.05 were correlated with nucleotides that were over or under-represented at a particular location.

**Figure 2.**
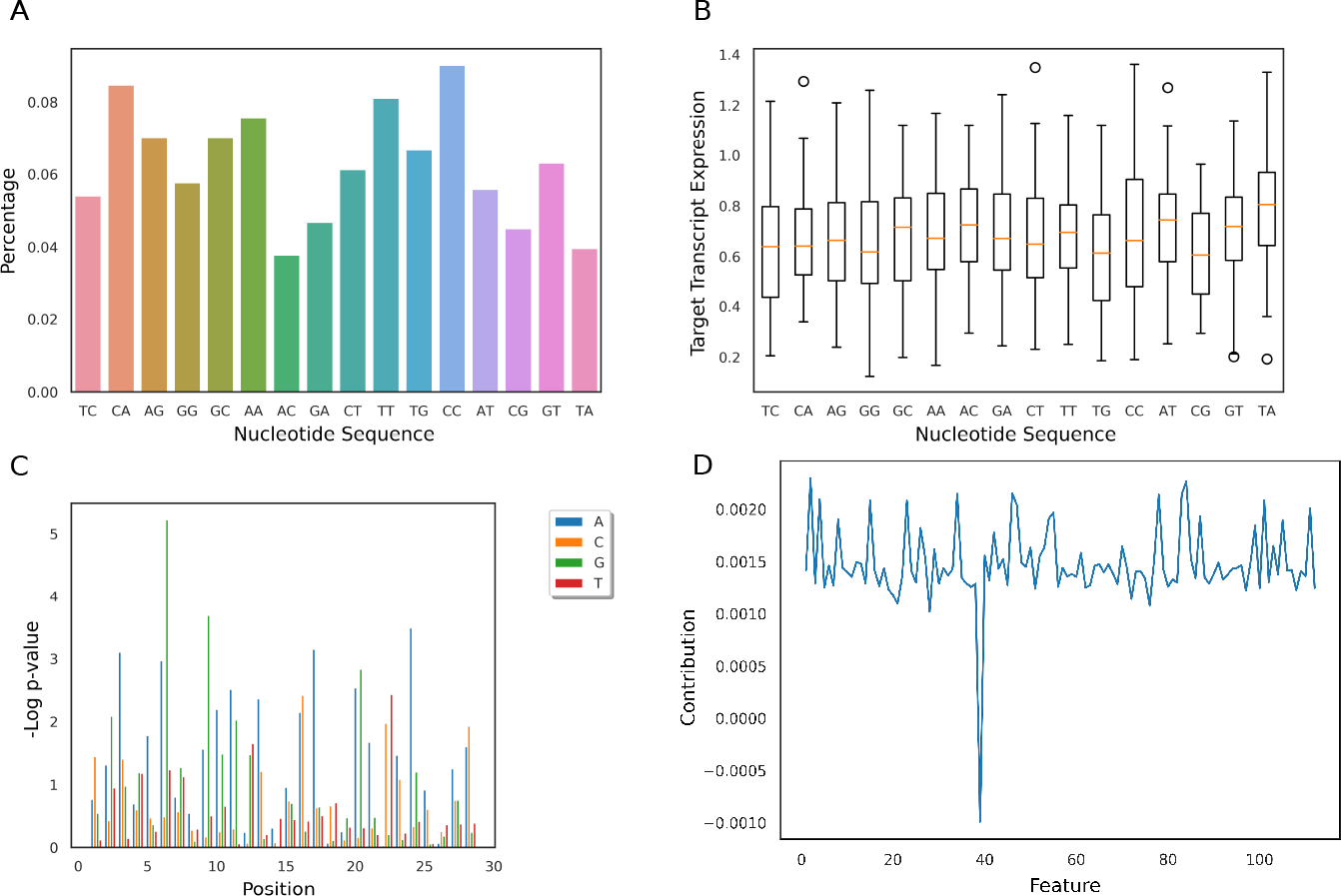
K-mer Analysis to study the guide composition. A: Bar plot of the population of sgRNAs that contain a specific dinucleotide at position 8. B: Box plot of target transcript expression values as a function of the nucleotide at position 8. C. Barplot of negative log of univariate linear regression significance p-value for all monomers at all positions across the guide. D: Bar plot of the feature contribution score for each feature in the Random Forest gini feature list.

The significance of each k-mer composition feature was then evaluated using the univariate linear regression module from sklearn. Next, all sequence features with p-values greater than 0.05 were removed (Fig 2C). Wary of being too inclusive with all statistically significant features, two additional subsets of these features were also considered, using a Z-score analysis on the negative log of p-values. Using 2 and 3 as the cutoff values, two sets of highly correlated statistical features were generated. In addition to this, a Gini score analysis was performed on each k-mer using the Decision Tree (18) and Random Forest (19) machine learning modules from sklearn. A similar approach was utilized by Fusi et al (20) for determining the most important features for CRISPR/Cas9 efficiency.

### Protein-RNA occupancy profile

In addition to the sequence composition features, we included transcriptome-wide occupancy as a feature into the model. To do so, we downloaded the transcriptome-wide protein occupancy data (raw reads in the FASTQ format) of HEK293 cells from Schueler et al (15). Reads were checked for adapter content and overall quality using Fastqc (21), trimmed using Trim Galore (22,23), and aligned to the human reference transcriptome (a combination of hg38 cDNA and ncRNA downloaded from Biomart (24) using hisat2 (25). After sequence alignment, peak calling was performed using macs2 (26), the resulting .xls file was used as an input for the model. Each guide sequence was also aligned to the human reference transcriptome, using tophat (27), allowing for 2 mismatches, the maximum number of mismatches tolerated by the CRISPR cas13 system (14). The indexed position of each guide on the target transcript was compared with the protein occupancy information for any overlap. The amount of overlap was recorded as a percentage of length of the guide and incorporated as a feature in the model.

### Composition features

In addition to the k-mer composition and occupancy features, a variety of other features were also included. These include guide spacer percent composition for each nucleic acid, guide location along the length of the transcript, and the number of complementary sequences in the reference transcriptome. Previous studies (28,29) have shown that RNA base composition plays a crucial role in not just the long term stability of the polynucleotide, but also in the activity of the Cas13 system. It is widely believed that the ends of transcripts, both 5’ and 3’ are highly structured, both to protect the transcript from degradation and to facilitate movement to different cellular compartments. This information was incorporated as a feature into the model, by calculating the midpoint of the complementary region of the transcript, and normalizing by the length of the target transcript. Despite the length of the guide, it is possible for guides to be complementary to different regions of different transcripts. This redundancy in targets, would possibly lead to off target effects, and significantly reduce the system’s ability to deplete a target transcript. To capture this in the model, the number of different hits returned from the tophat alignment for each guide was also recorded as a feature.

### Model architecture and feature selection

The occupancy and composition features, were then combined with the several sets of k-mer features (significant, Z-score > 2, Z-score > 3, decision tree Gini, and random forest Gini) and tested using a variety of machine learning algorithms, to determine the optimal combination. Each framework was evaluated based on their ability to accurately classify guides into one of four classes (0 – 3), based upon the quartile of transcript expression. This was tested by utilizing two different methods of cross-validation: 3-fold and 5-fold. For the 3-fold cross-validation, the experimental replicates were divided into separate folds, with two replicates serving as the testing data, and the other serving as an independent data set. For the 5-fold cross-validation the data from all 3 replicates were randomized, with 80% selected as training data and 20% selected for testing. The average values of the three different 3-fold experiments are presented in Table 1, as well as the average value of 100 different 5-fold experiments.

**Table 1.** Model Architecture Performance by Feature Set. Distribution of model accuracy using a variety of different architectures and different feature lists for both 5-fold and 3-fold cross validation methods. For KNN and Random Forest, average values for parameters with the highest accuracy are recorded in brackets.

| 3-Fold | P < 0.05 | Z > 2 | Z >3 | Gini | Decision Tree |
| --- | --- | --- | --- | --- | --- |
| Random Forest | 0.654 [41.3] | 0.651 [46.75] | 0.641 [45.75] | 0.652 [43.95] | 0.654 [41.3] |
| KNN | 0.558 [3.39] | 0.496 [9.71] | 0.515 [8.44] | 0.535 [3.67] | 0.558 [3.39] |
| SVC [linear] | 0.615 | 0.553 | 0.504 | 0.589 | 0.634 |
| **SVC [poly]** | **0.593** | **0.535** | **0.501** | **0.562** | **0.574** |
| **SVC [sigmoid]** | **0.551** | **0.489** | **0.458** | **0.517** | **0.521** |
| **SVC [rbf]** | **0.563** | **0.504** | **0.469** | **0.537** | **0.541** |
| **Decision Tree** | **0.634** | **0.636** | **0.634** | **0.640** | **0.637** |
| **5-Fold** | **P < 0.05** | **Z > 2** | **Z >3** | **Gini** | **Decision Tree** |
| **Random Forest** | **0.717 [55.4]** | **0.713 [57.55]** | **0.714 [51.1]** | **0.716 [54.35]** | **0.717 [50.55]** |
| **KNN** | **0.715 [2]** | **0.715 [2]** | **0.711 [2]** | **0.717 [2]** | **0.717 [3]** |
| **SVC [linear]** | **0.710** | **0.609** | **0.505** | **0.600** | **0.640** |
| **SVC [poly]** | **0.698** | **0.607** | **0.517** | **0.599** | **0.616** |
| **SVC [sigmoid]** | **0.620** | **0.551** | **0.470** | **0.540** | **0.553** |
| **SVC [rbf]** | **0.642** | **0.521** | **0.487** | **0.567** | **0.581** |
| **Decision Tree** | **0.715** | **0.715** | **0.711** | **0.715** | **0.714** |

Due to the experimental noise native to the data source methodology, a significant amount of the experimental replicates for a specific guide differed significantly in transcript expression, often by more than 25%. This discordance in the training data lead to the model receiving different labels for the same set of training features, imposing a hard cap to the model’s cross-validation performance. To account for this, the models were evaluated based upon noise-normalized accuracy. The noise-normalized maximum was calculated by counting the number of occurrences where one experimental replicate differed in transcript expression quartile, with another replicate of the same experiment. Put more formally by, computing the size of the set of tuples (i,j) such that x_i_ = x_j_ and y_i_ ≠y_j_, divided by the size of the set (i,j), and subtracting that value from 1 (where x and y correspond to the model input data and the label, respectively). The total model accuracy was then divided by the noise-normalized maximum to create the noise-normalized accuracy value.

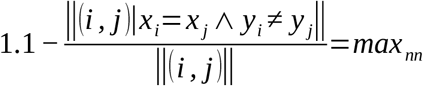

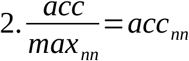

Once the optimal model architecture and feature set was determined, the importance of each feature was studied. To this end, the model was evaluated using 5-fold cross-validation 100 times, to establish a background. A single feature was removed from the model, and the model was evaluated another 100 times. The difference in model performance between the mean model accuracy and the background was taken to be the result of the removed feature (Fig 2D).

### POP-seq

Briefly, a total of 20 million cells were subjected to three variants of POP-seq including UV crosslinking, Formaldehyde crosslinking and No-crosslinking approaches (as described in Srivatava et al. (30)). Cells were lysed in trizol and the resulting interphase layer was treated with RNase A/T1, Proteinase K, DNase I and depletion of r-RNA. 50 ng of high quality RNA was used to prepare libraries for small RNA Illumina sequencing.

RNA purity and concentration were assessed at each step using Nanodrop, based on the absorbance ratio 260/280 >2. RNA integrity was evaluated using Agilent 2100 bioanalyser system. Atleast 50 ng of r-RNA depleted RNA was used to generate sequencing libraries using the True-seq small RNA library prep kit (Illumina). All libraries were barcoded and sequenced in parallel on a Next-seq platform for 400 million reads to obtain 75 bp single end reads.

## Results

CASowary takes a list of gene names as input and exports a list of sgRNA sequences predicted to be at least efficient, with a transcript expression value between 0.5 and 0. The tool first collects a list of Ensembl transcripts that map to the input genes, then creates all possible 28 nucleotide guides that span the length of those transcripts and saves them in a FASTA file. The FASTA file is then aligned to the reference transcriptome using tophat, allowing for 2 mismatches, to create a BAM file. The resulting BAM file is converted to a BED file using bedtools (31). That BED file is then fed into the model where it classifies each guide; and outputs a separate text file for each transcript mapping to an input gene name, containing all highly effective guide sequences ranked upon model confidence in its classification.

Our tool uses a Decision Tree architecture and set of features based upon Random Forest Gini analysis to classify a sgRNA into 1 of 4 classes, based upon predicted transcript knockdown efficiency. Each class represents a specific quartile of normalized expression (0: 0 – 0.25, 1: 0.25 – 0.5, 2: 0.5 – 0.75, and 3: 0.75 – 1). Guides belonging to class 0 and 1 were categorized as highly efficient and efficient, while classes 2 and 3 correspond to inefficient and highly inefficient, respectively. Utilizing 5-fold and 3-fold cross-validation, this model was benchmarked with noise-normalized accuracy of 70.1% and 74.3% respectively (69.2% and 71.5% without accounting for noise in the source data). A small amount of overfitting was observed in the 3-fold cross-validation, due to the identical model inputs, so 70.1% was believed to be the most accurate measure of the model’s performance.

Using one class vs all pairwise comparisons, a Receiver Operating Characteristic (ROC) curve for the model was created (Fig 3). Calculating the Area Under the Curve (AUC) for each class revealed that the model performed best predictions for highly efficient and highly inefficient guides (0: 0.949, 1: 0.869, 2: 0.753, and 3: 0.839). These numbers clearly illustrate an increased sensitivity in model’s predictions for highly efficient and efficient class of guides, maximizing its effectiveness.

**Figure 3.**
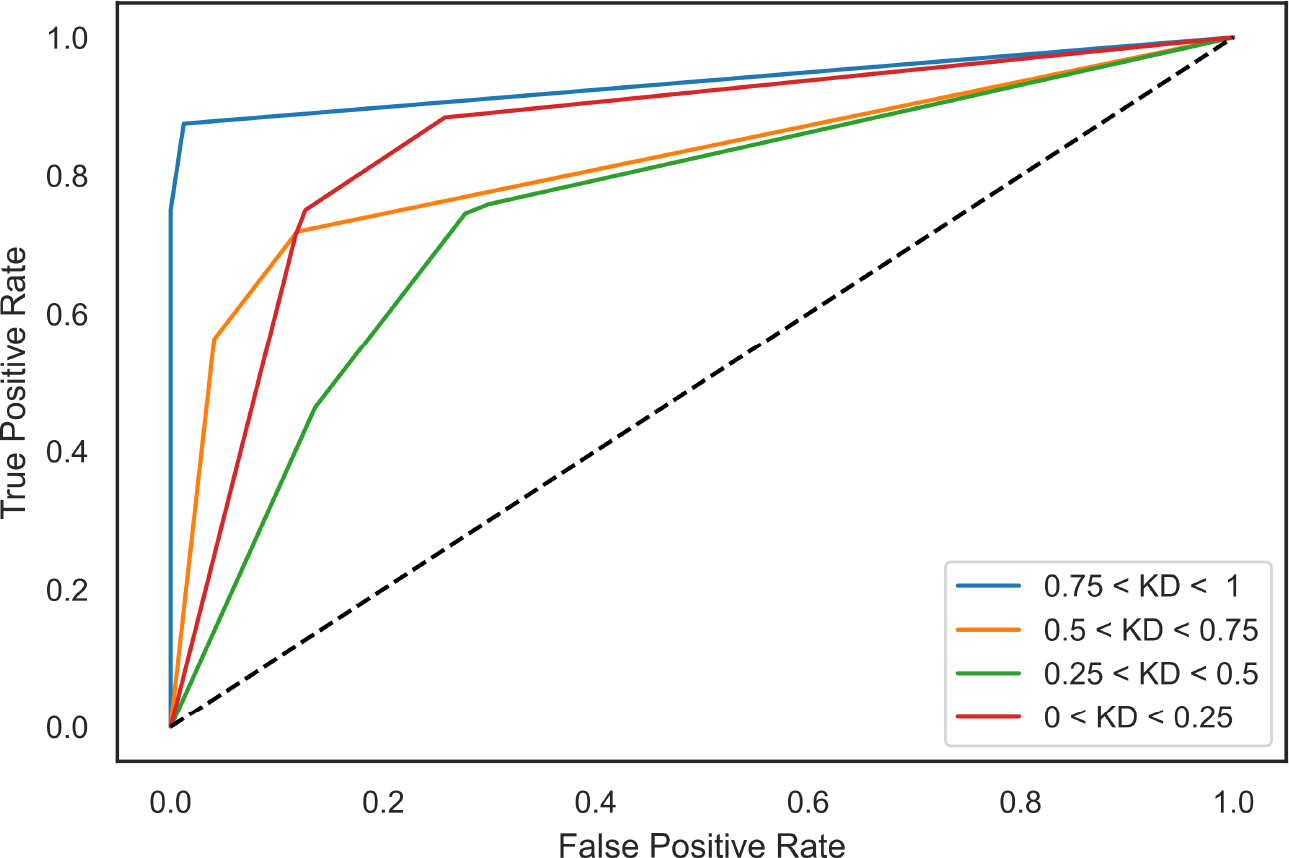
CASowary Model Performance. ROC curve for CASowary Decision Tree model using Random Forest feature list.

We performed a predictive analysis against 5 different genes, including the SMARCA4 gene. We conducted a series of characterization experiments using CIRTS (CRISPR-Cas-inspired RNA targeting system), an orthogonal RNA-targeting system (16,32). A series of guides targeting a single SMARCA4 transcript (ENST00000344626.9) were obtained from IDT and the transcript depletion experiment data was generated and analyzed for a select set of sgRNA predictions (Fig 4) (16). Comparing our model’s predictions of high (best) and low (worst) efficiency guides and the experimental results of CIRTS showed a very high correlation (Fig 4). This experimental confirmation illustrates the robust and reproducible nature of predictions made by CASowary using an orthogonal RNA editing system to Cas13.

**Figure 4.**
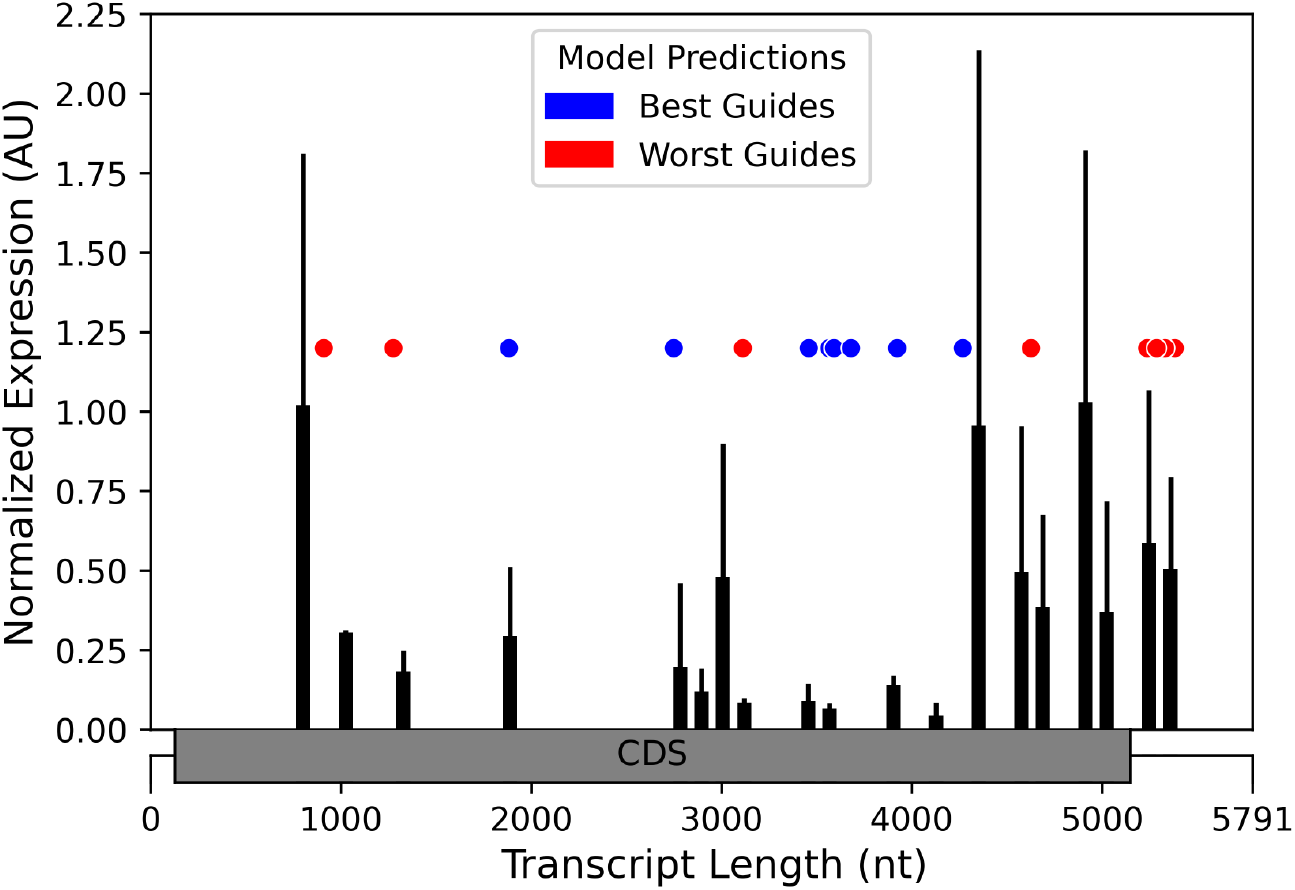
Comparison of CASowary Predictions with CIRTS Results. CIRTS experiments SMARCA4 (add transcript ID) transcript measurements correlated with high efficiency CASowary guide predictions, transcript expression value between 0.25 – 0 (red) and low efficiency CASowary guide predictions, transcript expression value between 0.75 and 1 (blue).

During the development of CASowary, we observed that specific classes of guides exhibited preferential patterns across the length of the transcripts. To confirm this trend, a comprehensive analysis of the predictions for a random assortment of 5900 gene transcripts were performed, resulting in 12.7 million mapped guides. All guides of a specific class were then grouped and plotted against their corresponding location in the transcript, from 5’ to 3’ direction, by normalizing the length to understand locational preferences for various classes of guides. This analysis revealed that the majority of the guides were predicted to be inefficient, either categorized as Highly Inefficient or Inefficient (90.7%) (Fig 5 C-D). In addition, our data suggests that efficient guides (Efficient and Highly Efficient) primarily reside in the intermediate regions of the transcript, especially between 30% and 70% the length of the transcript (Fig 5A-B). Distribution of the guide locations was similar when we plotted the data for the complete training data (Fig 5A) as well as the computational guide predictions for 5900 genes (Fig 5B). This observation supports the theory that the ends of active mRNAs are highly structured, with multiple binding proteins and other modifications that further limit the binding efficiency of the CRISPR-Cas13 system.

**Figure 5.**
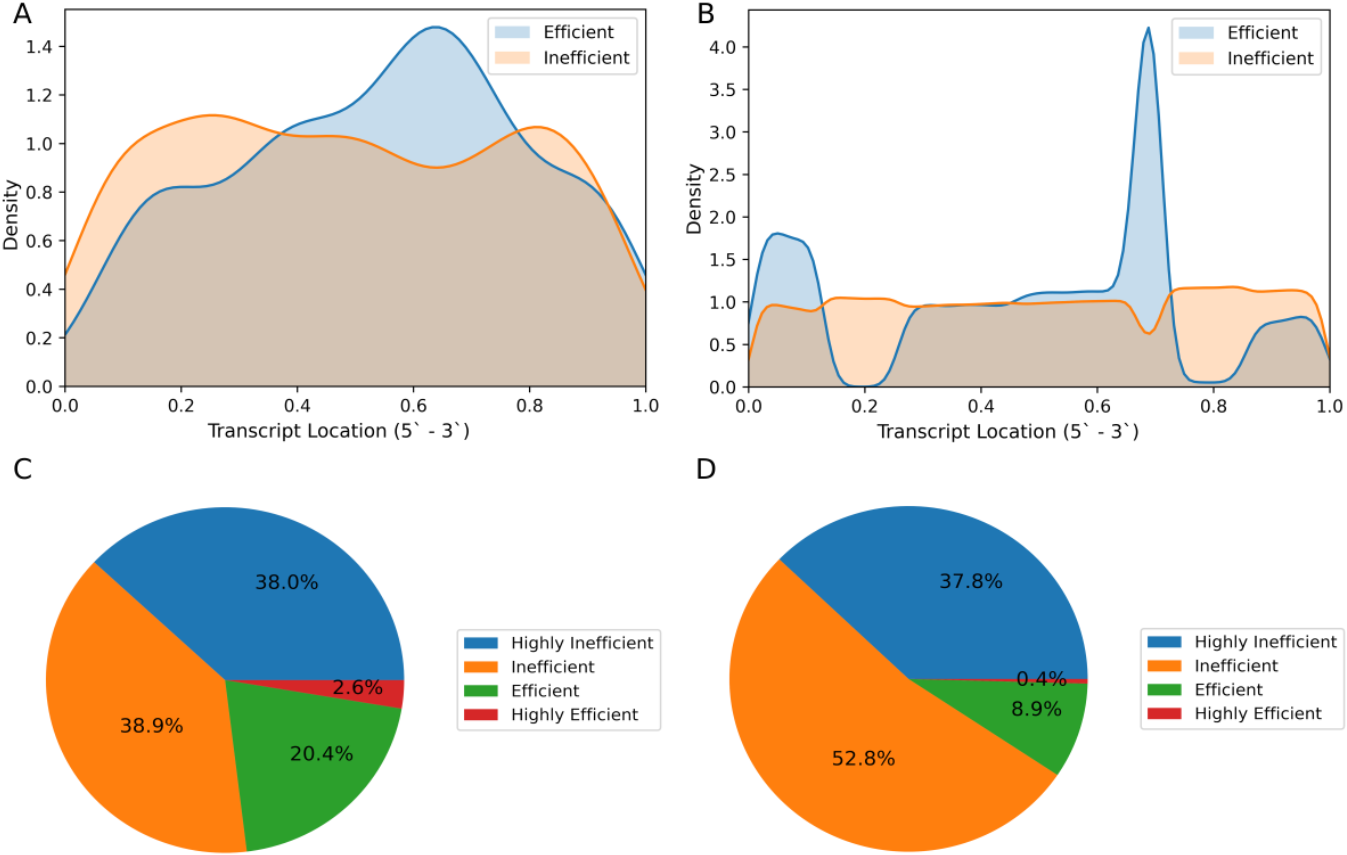
Comparison of Training Data with Gene Predictions: A: Density plot of Efficient (Highly Efficient and Efficient) and Inefficient (Inefficient and Highly Inefficient) guides from the training data. B: Density plot of Efficient and Inefficient guides from the 5900 random genes. C: Pie chart for the breakdown of guide predictions from the training data. D: Pie chart for the breakdown of the guide predictions from the 5900 random genes.

The secondary location for efficient guides, lying between 0% and 20% of the transcript (near the 5’ end) among the computational predictions (Fig 5B), was unexpected and required additional investigation. Of the 5900 genes included in the analysis, transcripts from 4361 different genes included efficient guides in the upper quintet. That subset included 1417 different genes associated with lncRNA (32%), over 90% of all genes (1570) associated with lncRNA. The average length of these transcripts (1670 nucleotides) was significantly longer than the average length of all transcripts (1537 nucleotides) with p-values from Mann-Whitney (33) of 1.34 × 10 ^-43^. This illustrates that longer transcripts are more likely to have guides in this region, and that this region is the prime target for lncRNA depletion.

Next, we investigated the ability of CASowary to generate cell type specific guide predictions by employing the tool to predict guide sequences on the HeLa cell line. To this end, we utilized in-house phase separation based protein occupancy data for the HeLa cell line, through a method called Protein-Occupancy Profile Sequencing (POP-seq) to map protein-RNA interactions on a transcriptome wide scale (30). Protein RNA-interactions are known to vary from cell type to cell type, which would alter the accessibility feature of the model (34,35).

The reads from the experiment, corresponding to stretches of RNA molecules interacting with proteins, were run through the same computational pipeline as described in methods. The resulting file was substituted for the HEK293 peak file in CASowary’s input. A list of 100 candidate genes with differential binding profiles between the HEK293 and the HeLa files was generated by running them through DiffHunter (36). This list of candidate genes was then analyzed using CASowary with the HeLa occupancy profile. Transcript levels were verified by comparing prevalence of reads supporting a specific transcript from RNA-Seq experiments on the wild type of that cell line. This data was obtained for both HEK293 and HeLa cells from the Gene Expression Omnibus (GEO) (37), series accession number GSE146946.

The results of CASowary predictions for the two cell lines were visualized using Integrative Genomics Viewer (38) along with their relative abundance (SRR11304482 and SRR11304484, for HEK293 and HeLa respectively) (39). Comparing the two cell line results revealed high correlation for guides outside of protein occupied regions. However, regions with differential protein binding exhibited a significantly altered pattern of predicted guides across the transcript (Fig 6). The results from the analysis indicate that CASowary’s guide predictions are highly sensitive to protein occupancy, allowing for cell type specific guide predictions. This exciting development illustrates the importance and need for more in-depth protein occupancy protocols to enable guide predictions on gene regulatory regions tailored for specific tissues and cell types.

**Figure 6.**
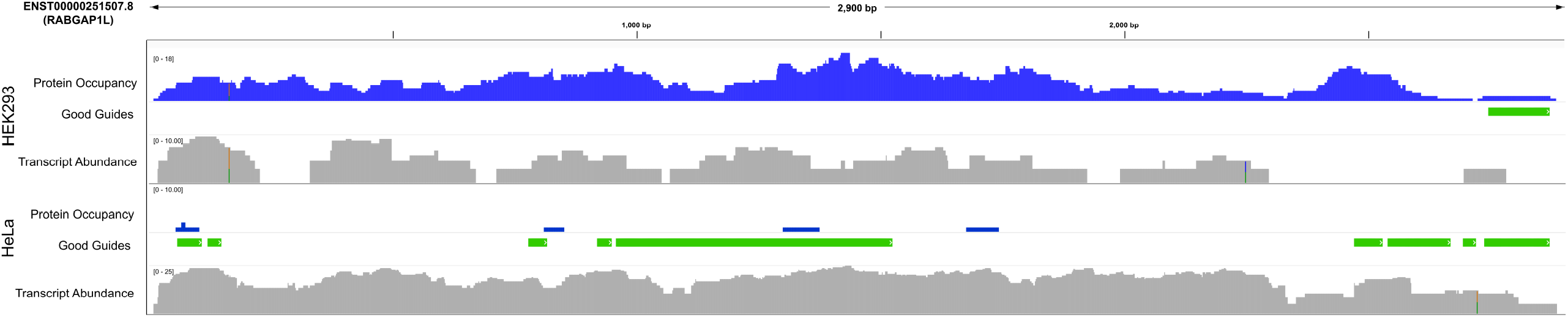
Cell Line Specific Predictions: IGV tracks for ENST00000554738.5, a protein coding transcript for NRXN3. The top collection of tracks corresponds to protein occupancy (blue), high quality guide locations (transcript expression value between 0 and 0.5) (green), and transcript abundance for HEK293 cell line (gray). The bottom collection of tracks is the same for the HeLa cell line.

## Conclusions

Gene and transcript editing technologies, such as CRISPR and its variant systems, will continue to evolve for their application, and so too will the demand for computational and predictive tools to improve the efficacy of these methods. We present CASowary as the first of its kind, tool that provides RNA targeting CRISPR support software. Utilizing the selective set of sequence and RNA accessibility features, our tool can generate a list of potential sgRNAs predicted to be highly efficient, from among thousands of possible guides. Therefore, CASowary’s predictions open the door for new RNA based gene therapies and personalized medicine.

Despite the success of the current iteration of our tool, there still remains room for improvement. In the future, we aim to incorporate additional availability information by considering the structure of the target RNA *in vivo*. There is also a desire to expand the cell and tissue specific predictions, but that requires substantially more protein occupancy information.

### Availability and requirements

CASowary is written in Python, requiring 3.6.8 or above, with some dependencies on Python 2.7.16. Source code for CASowary is available for free for academic use under GitHub (https://github.com/Janga-Lab/CASowary).

## Abbreviations

AUC: Area Under the Curve
Cas: CRISPR associated
cDNA: complementary DNA
CIRTS: CRISPR-Cas-Inspired RNA Targeting System
CRISPR: Clustered Regularly Interspersed Short Palindromic Repeats
crRNA: crispr RNA
DNA: DeoxyRibonucleic Acid
GEO: Gene Expression Omnibus
IDT: Integrated DNA Technologies
IGV: Integrative Genomics Viewer
lncRNA: long non-coding RNA
ncRNA: non-coding RNA
NIH: National Institute of Health
PAM: Protospacer Adjacent Motif
POP-seq: Protein Occupancy Profile sequencing
RBP: RNA Binding Protien
ROC: Receiver Operator Curve
sgRNA: single guide RNA

## Declarations

### Ethics approval and consent to participate

NA

### Consent to publish

NA

## Availability of data and materials

All Pop-seq data generated in this study is deposited under GEO accession number GSE166189. Reviewers can confidentially access the data via https://www.ncbi.nlm.nih.gov/geo/query/acc.cgi?acc=GSE166189 and entering token ‘mtybqumuzxuzrkz’ until the data is made open access.

## Author’s Contributions

AK wrote the code base, performed the computational benchmarking, and wrote the manuscript. MS performed the laboratory work for POP-seq data generation. SR performed CIRTS experiments to validate guide RNA efficacy. RS analyzed POP-seq data. BCD supervised the CIRTS characterization experiments. SCJ oversaw the creation of the tool, collaboration with external colleagues, and contributed to drafting the manuscript. All authors have read and approved the final manuscript.

## Acknowledgments

The authors would like to thank the respective labs of SCJ and BCD for their advice and support over the course of the project. AK, MS, and SCJ would like to thank Dr. Mark Kaplan for providing space and access to equipment for wet lab experiments.

